# Probing How Anti-Huntingtin Antibodies Bind the Fibrillar Fuzzy Coat Using Solid-State NMR

**DOI:** 10.1101/2025.04.25.650083

**Authors:** Raffaella Parlato, Greeshma Jain, Alessia Lasorsa, Patrick C.A. van der Wel

## Abstract

Antibodies are critical for the immune response and serve as important tools due to their ability to recognize specific amino acid sequences, or epitopes. Based on the latter, they are utilized as diagnostic tools in biological and biomedical research. Huntington’s disease (HD) is a neurodegenerative condition caused by CAG repeat expansions in the huntingtin (HTT) gene and characterized by amyloid-like protein deposits in patients. Multiple anti-HTT antibodies are used in HD research for their ability to recognize specific HTT inclusions in both post-mortem tissue and in laboratory conditions. Some of the antibodies are seen as detectors of distinct structural motifs. However, most knowledge of their binding mechanism stems from studies of soluble monomers or short fragments of the epitopes, rather than the aggregated, misfolded target protein. Here, we investigate how MW8 antibodies interact with HTT exon 1 (HTTex1) fibrils, using solid-state NMR, electron microscopy, and complementary techniques. Magic angle spinning NMR revealed localized impacts of the antibody on exposed parts of the HTTex1 fibrils: the flanking segments that form its ‘fuzzy coat’. Antibody binding affected the structure and dynamics of the fuzzy coat, but also modulated the propensity for forming supramolecular fibril clusters, which has important implications for (reducing) cytotoxicity.

## Introduction

Amyloid diseases encompass a group of disorders characterized by the abnormal accumulation of misfolded proteins in various tissues and organs, with notable examples including Parkinson’s disease, Alzheimer’s disease, and Huntington’s disease (HD). HD is an incurable autosomal dominant neurodegenerative disorder characterized by involuntary movements, cognitive decline, and psychiatric symptoms. This condition arises from a mutation involving the expansion of an uninterrupted CAG repeat, encoding for a stretch of glutamines (polyglutamine or polyQ) in the huntingtin (HTT) protein.^[1]^ Expansion beyond a threshold of 35 codon repeats triggers the disease, with a longer CAG repeat stretch correlating with an earlier age of onset. In patient cells, a hallmark of HD pathology is the formation of mutated HTT exon 1 (HTTex1) fragments,^[2,3]^ which are highly aggregation-prone, interfere with the cellular pathways, and may contribute to disease progression.^[4]^ Given the pathological significance of HTTex1 generation, misfolding, and aggregation, HTTex1-binding antibodies have emerged as indispensable tools for both research and potential therapeutic approaches.^[5,6]^ DiFiglia et al. used anti-HTTex1 antibodies to show that the HTT aggregates derived from patient materials contained N-terminal fragments of mutated HTT.^[7]^ HTT-binding antibodies have been deployed in research on both the full-length HTT protein and various HTT fragments. A key development has been the production of monoclonal antibodies, such as MW1 and MW8, that specifically target sub-domains of HTTex1: the N-terminal segment (HTT^NT^), the polyQ segment (which contains the mutation and forms the core of the fibril), and the C-terminal proline-rich domain (PRD).^[8–10]^ The latter, together with HTT^NT^, forms the fuzzy coat of HTTex1 aggregates, as illustrated in our recently published integrative structure of HTTex1 fibrils (Figure 1a).^[11]^ The MW1 antibodies bind to the polyQ segment, while MW8 antibodies recognize epitopes located at the very C-terminus of the PRD (Figure 1a-c).^[8]^ Such antibodies have proven to be invaluable tools for characterizing protein aggregates, as they can discriminate between different conformational states, aggregation stages, and polymorphisms. Crucially, epitope accessibility across the three HTT domains varies during the fibril formation process, and also among mature aggregates due to aggregate polymorphism, as explained below.^[8,12– 15]^ MW1 has low affinity for fibrils in which the polyQ segment is buried, while MW8 can still bind exposed epitopes on the surface of the same fibrils.^[13,15]^ Antibodies can interact with epitopes in HTTex1 fragments at various stages of aggregation, providing mechanistic insight and potentially reducing the pool of aggregation-prone fragments.^[5,7]^ Thus, beyond their role in detecting and characterizing aggregates, antibodies are being investigated as modulators of protein aggregation. In a therapeutic context, antibodies have recently shown promise in other amyloid diseases, such as Alzheimer’s disease, by targeting the aggregated species.^[16]^ It has been proposed that one can thus slow down the disease progression by reducing the Aβ monomer and oligomer pool,^[5,16,17]^ and by enhancing amyloid clearance.^[18]^

**Figure 1.**
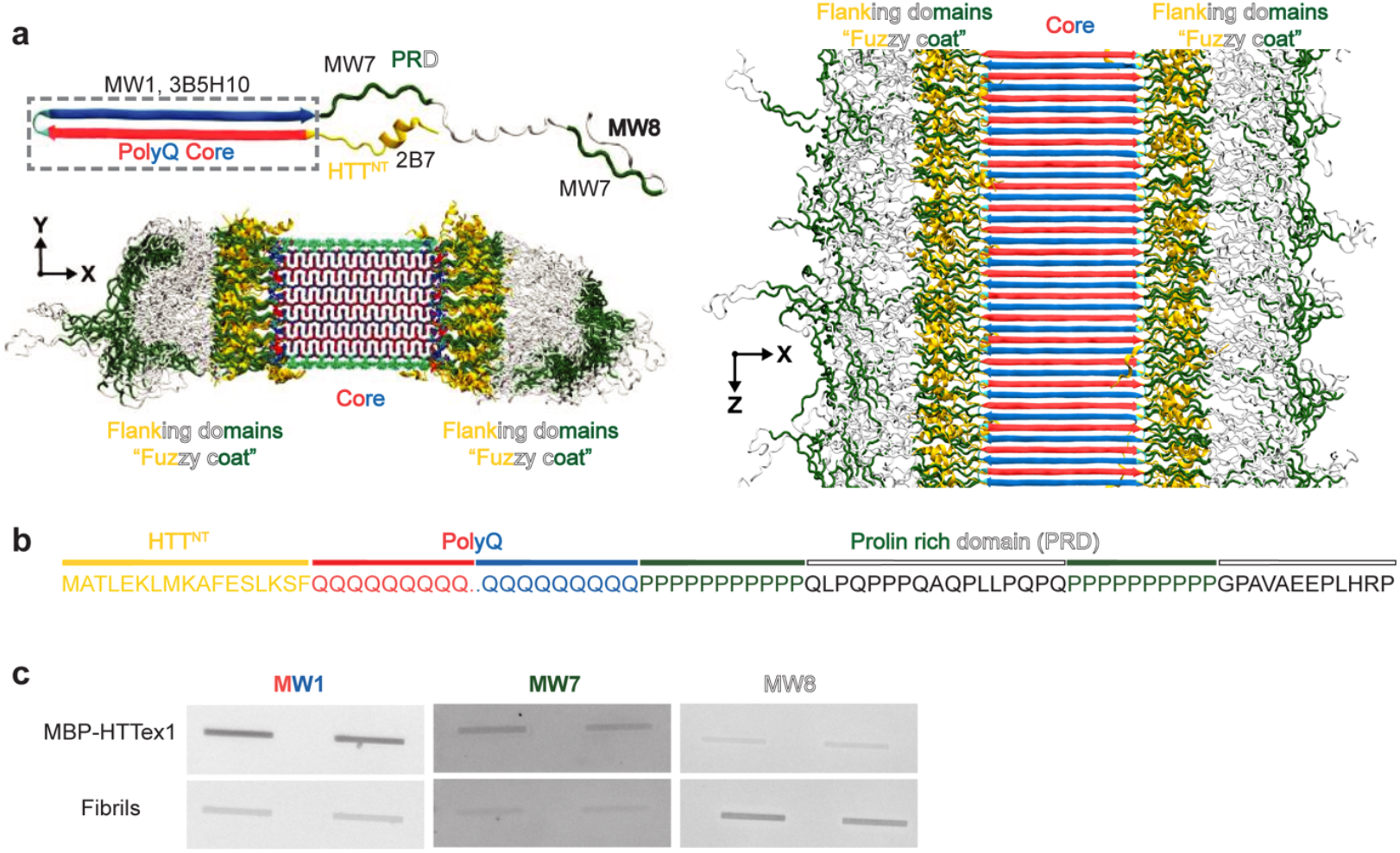
Structural overview of HTTex1 and its interaction with antibodies. a) Fibrillar structure of Q44-HTTex1 shown for the monomer (top) and intact fibrils (bottom and right). Three domains are indicated: HTT^NT^ in yellow, polyQ in red and blue, and PRD in green and white. The dotted box marks the monomer part sequestered in the fibril core. For each HTTex1 region, selected relevant antibodies are indicated (top left). b) HTTex1 primary sequence with the three domains named and color-coded as per panel a. c) Dot blot analysis of Q32-HTTex1 fibrils in monomeric (fusion protein MBP-HTTex1) and fibrillar states, using the antibodies MW1, MW7, and MW8. The antibodies are colored according to the binding region. Panel a was adapted from reference ^[11]^ under its creative commons license.

The accessibility of epitopes relates to the burial and exposure of different domains of HTTex1. As noted (Figure 1), HTTex1 fibrils feature a fuzzy coat formed by the non-polyQ protein segments. Fuzzy coats are common structural features of pathogenic amyloid fibrils and are of general interest. For example, the tau fuzzy coat can engage in interactions with other biomolecules, affecting the production of soluble and toxic species.^[19]^ The Ab fibril fuzzy coat is the point of interaction for chaperones,^[20]^ which recognize regions in the flanking regions, influencing both the aggregation and disaggregation pathways.^[21]^ Aside from these interactions, exposed flanking regions may be recognized by ubiquitin ligases, which facilitate protein clearance through the ubiquitin-proteasome system. The dysregulation of this pathway compromises the removal of amyloid and is implicated in the pathogenesis of HD and other amyloid disorders.^[22]^ However, the study of fuzzy coats and their interactions remains challenging due to the intrinsic structural flexibility and heterogeneity (Figure 1a). The flanking domains adopt multiple conformations even within a single fibril species and exhibit extensive dynamics behavior. As a result, high-resolution structural techniques like cryo-electron microscopy and X-ray crystallography often struggle to resolve their structure.

A key concept for these types of antibodies and their applications relates to the idea that they can specifically or selectively recognize distinct structural or conformational states.^[13,14]^ However, whether or how this works remains incompletely understood on the molecular level, emphasizing the need for further research. A detailed understanding of the structural interactions between antibodies and their amyloid targets is essential for advancing antibody-based therapies. Structural insights at atomic resolution are critical for rational antibody design, enabling the optimization of binding affinity, selectivity, and therapeutic efficacy^[23]^. Importantly, such insights can reveal how antibodies could discriminate between pathogenic and non-pathogenic protein conformations, a key consideration given the structural heterogeneity of amyloid aggregates.

Until now, most structural data on HD-relevant antibodies have been derived from X-ray crystallography studies.^[24– 28]^ However, such methods are limited to an analysis of the antibody without substrate, or with short peptide fragments or protein monomers. They do not shed light on the interaction with actual misfolded protein aggregates. In prior work, we and others have deployed magic angle spinning (MAS) solid-state NMR (ssNMR) spectroscopy as a powerful tool to probe HTTex1 fibrils.^[11,29–31]^ This enabled an atomic-level description of the fibrils (Figure 1a) via an integrative structural biology approach.^[11]^ Fibrils consist of individual HTTex1 monomers that assemble into a central rigid core surrounded by flexible domains. Within the core, the polyQ domain adopts a β-sheet conformation (red and blue arrows) (Figure 1a), outside of which dynamic flanking domains form the fuzzy coat.^[11]^ In this study, we deployed analogous ssNMR and complementary structural techniques to probe how MW8 antibodies bind HTTex1 fibrils. We observed the effect of binding on protein mobility and structure at the level of protofilaments, and also examined the impact on fibril clustering. These studies combine ssNMR, fluorescence-based assays, and transmission electron microscopy (TEM) to provide a comprehensive analysis on different length scales.

## Results and Discussion

### Anti-HTT Antibody MW8 Binding to HTTex1 Fibrils

A variety of anti-HTTex1 antibodies are available to investigate HTTex1 and its fibrillar state.^[8–10,32–38]^ We performed immunoblotting studies with select antibodies designed to target different epitopes in the HTTex1 domains: MW1 targets the polyQ domain, MW7 recognizes oligo-proline sequences (found in the PRD), and MW8 binds sequence AEEPLHRP, in the very end of the PRD (Figure 1a, b).^[8,13,15]^ HTTex1 fibrils were formed from Q32-HTTex1 proteins, via spontaneous aggregation following previously established conditions (in the absence of antibody).^[39]^ As a monomeric control, we also used a fusion protein of HTTex1 attached to the 42.5 kDa-sized maltose binding protein (MBP), which prevents aggregation.^[40,41]^ Using fluorescently labeled secondary antibodies, we observed distinct differences in fluorescence signal upon antibody binding to these fibrillar and monomeric states of HTTex1. MW1 and MW7 showed a stronger affinity for monomers; in contrast, MW8 antibodies exhibited strong binding to HTTex1 fibrils (Figure 1c).

The antibody 2B7 showed similar results to MW8 (Figure S1). These observations are consistent with their epitopes being outside the fibril core in the exposed flanking segments (Figure 1), and are in line with prior reports.^[13,15]^ Seeing a reduced fluorescence signal for fibrils suggests that the epitope is buried in the fibrillar state (e.g., the polyQ segment detected by MW1). In contrast, the strong binding of MW8 to its epitope in the fibrillar state supported the idea that its epitopes are exposed in the fibrils. Our data are consistent with prior reports of strong MW8 binding to HTTex1 fibrils seen *in vitro*, in cells, and in patient materials.^[5,8,13,15,31]^ Indeed, MW8 has turned into a commonly used diagnostic tool in HD research.^[42]^ Notably, MW8 recognizes the very end of the PRD domain, which, in the full-length protein, is not available for antibody recognition due to the steric hindrance of folded HEAT repeat domains (see cryo-EM structure in Figure S2a-c).^[43,44]^ In our model of the HTTex1 fibril, the MW8 epitope is accessible, permitting the recognition of HTTex1 fragments (Figure S2d).^[8,11,31]^ Because of this distinction between full-length HTT and aggregated HTT fragments, MW8 has been employed as an aggregate detector in high-sensitivity assays.^[42,45,46]^

### Impact of MW8 on Supramolecular Fibril Architecture

Next, we probed structurally how the antibody was interacting with the fibrils and whether it was altering the fibril architecture and morphology. First, negative stain TEM was performed on Q44-(Figure 2a and S3) and Q32-HTTex1 fibrils (Figure S4), MW8 alone (Figure 2b and S5), and Q44-HTTex1 fibrils bound with MW8 antibody, at molar ratio of 4:1 (monomeric HTTex1: MW8) (Figures 2c, S6). In the absence of antibodies (Figure 2a), the HTTex1 fibrils exhibited their typical bundling and filamentous morphology, with a diameter ranging from ~6 to ~12nm (Figure S3g).^[15]^ The antibodies alone (Figure 2b) appeared globular in shape, with a size comparable to their expected diameter of ~10-12 nm (yellow dot). A PDB file of a structurally similar monoclonal antibody from the IgG2a family was analyzed in ChimeraX, to provide an on-scale model alongside Q44-HTTex1 fibrils (~6nm core) (Figure 2d).^[11]^ TEM micrographs of fibrils incubated with MW8 (Figure 2c) revealed that the fibrils maintained their fibrillar shape, but they appeared surrounded by spherical objects with a globular shape, comparable in size to that expected for the antibodies (yellow dot) (Figure 2c, d).^[50]^ HTTex1 fibrils are well known to exhibit a characteristic tendency to bundle, clustering together to form larger supramolecular structures (both in cells and *in vitro*).^[51,52]^ Interestingly, in the presence of MW8, the HTTex1 fibrils appeared more bundled, and the possibility of finding isolated fibrils decreased compared to the control (Q44-HTTex1 alone). The bundling became more pronounced, leading to a decreased occurrence of isolated fibrils. The large clusters formed upon the antibody-fibril interaction suggest that the antibodies bridge epitopes on different fibrils, leading to more extensive fibril bundling. Notably, bigger inclusions are generally thought to be less toxic than smaller aggregated states (e.g., individual filaments, oligomers, and protofibrils)^[53,54]^.

**Figure 2.**
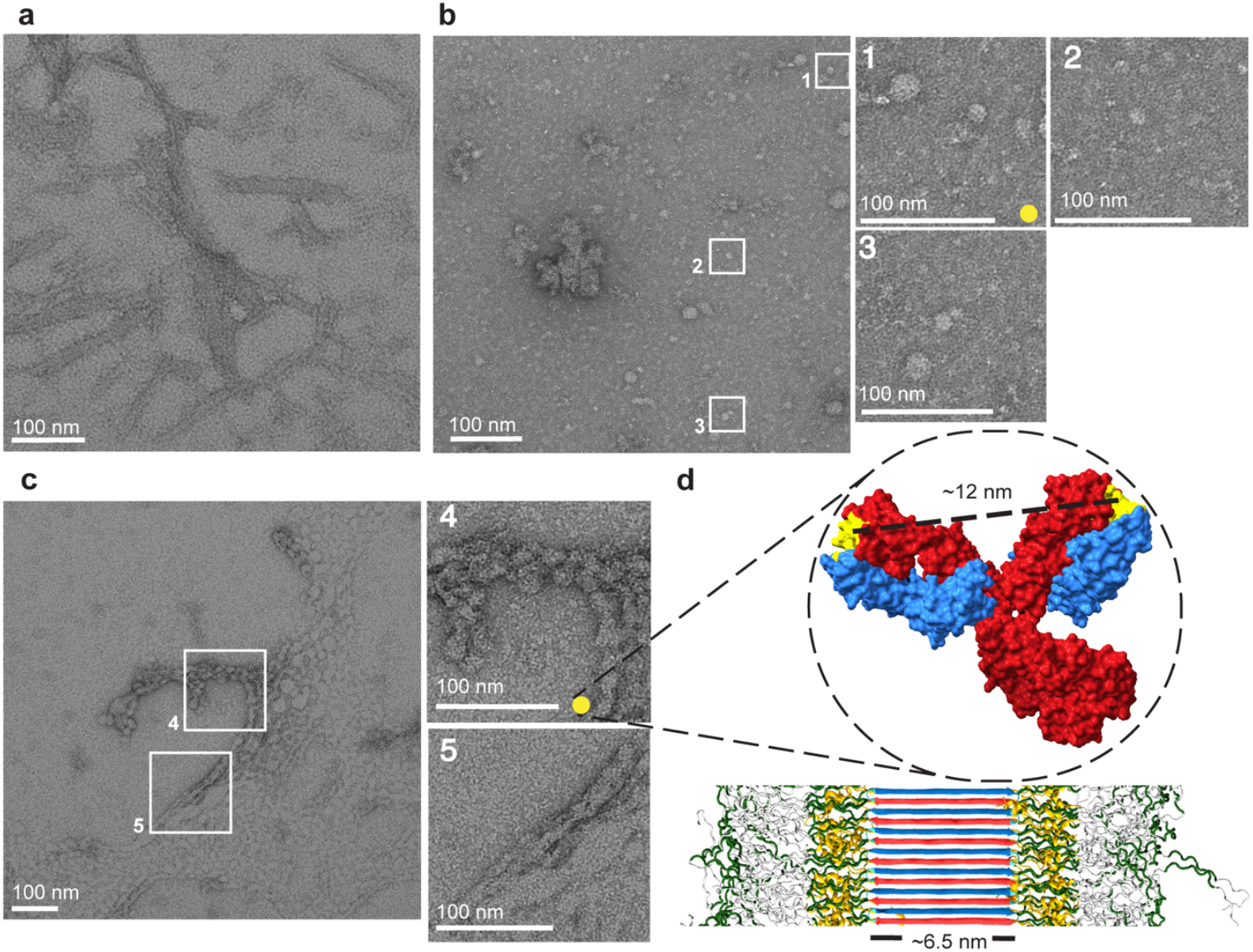
Negative stain TEM micrographs of HTTex1 fibrils and antibodies. a) Q44 HTTex1 fibrils. b) MW8 antibody alone; zoomed regions (white boxes 1, 2, and 3) illustrate the dimension of a single antibody (~12 nm, yellow dot). c) HTTex1 fibrils incubated with MW8 antibodies. In the white boxes 4 and 5, a zoomed region of the sample. d) On-scale fibril model indicating the polyQ core (~6.5nm width) and an IgG antibody (PDB: 5DK3)^[47]^. The antibody light and heavy chains are colored blue and red, respectively; binding sites are yellow. The models indicate the respective sizes of IgG antibodies and HTTex1 fibrils. The distance between the two binding sites is ~12nm, calculated using UCSF ChimeraX.^[48,49]^ The fibril model was adapted from reference ^[11]^ under its Creative Commons license.

### MAS NMR Analysis Shows No Changes in Fibril Core

To investigate the MW8-HTTex1 interactions at the molecular level, ssNMR was performed using ^13^C,^15^N-labeled HTTex1 fibrils. HTTex1 fibrils were prepared from isotope-labeled proteins in the absence of antibody. The resulting fibrils were split into two portions and used to prepare NMR samples. One sample of ~3.7 mg was prepared without antibodies and used as a control. Another portion of ~0.1 mg was incubated with the MW8 antibody at a 4:1 molar ratio overnight, prior to the preparation of the ssNMR sample. The limited sample size reflected a limiting amount of the MW8 antibody. First, ssNMR was used to determine whether MW8 binding affects the core of the fibril, which is constituted by the polyQ segments (Figure 1). To examine the structural effects on the core, cross-polarization (CP) ssNMR experiments were conducted on both samples (Figure 3a; Figure S7-S8). ^13^C CP ssNMR experiments suppress signals from mobile residues and enhance those from rigid residues, such as the polyQ core (Figure 3b, red/orange region).^[12,29,40,55]^ The resulting ^13^C 1D spectra resemble those previously seen for HTTex1, allowing the use of known assignments (Figure 3a). Two sets of peaks are seen in the C_α_ (~54 and 56 ppm) and C_β_ region (between 30 and 34 ppm), corresponding to two conformational states of glutamines within the core. The two conformations, designated as types “a” and “b”, are present in approximately equal amounts. Previous work has shown that this reflects two glutamine conformers that are forming alternating β-strands in the polyglutamine antiparallel β-sheet structure.^[30,39]^ Additionally, peaks corresponding to rigid and semirigid proline regions, P_A_ and P_B,_ are visible. As shown by the overlay of the HTTex1 and HTTex1+MW8 spectra in Fig. 1a, no significant changes in the fibril core signals were observed upon MW8 binding, suggesting that the core is not significantly affected by MW8 binding.

**Figure 3.**
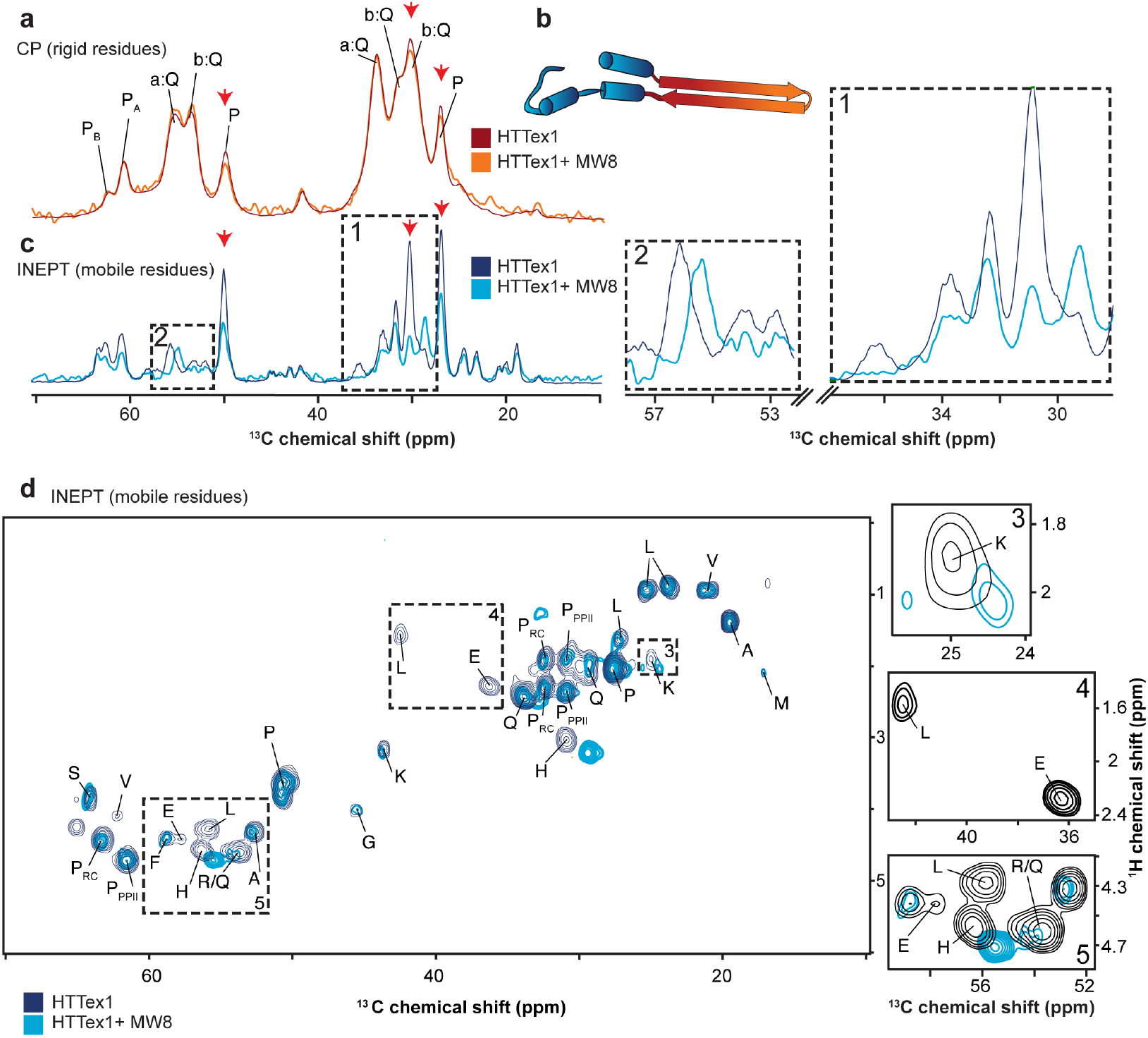
1D and 2D MAS ssNMR on labeled HTTex1 fibrils with and without MW8 antibody. a) 1D ^13^C CP spectra of HTTex1 (dark red) and HTTex1+MW8 (orange) show the rigid and semirigid residues from the fibril core. b) Structural representation of a single aggregated HTTex1: the polyQ domain in β-hairpin conformation (orange/red), and the exposed HTT^NT^ and the PRD (dark/light blue). c) 1D ^13^C INEPT spectra show signals from the flanking domains of HTTex1 (dark blue) and HTTex1+MW8 (light blue). The zoomed regions (dashed boxes 1 and 2) emphasize key differences. The red arrows indicate the intensity reduction in the 1D spectra. d) 2D INEPT spectra of HTTex1 fibrils (dark blue) and HTTex1+MW8 (light blue), with each peak labeled by the amino acid. Dashed boxes 3, 4, and 5 mark significant differences. The color coding in the spectra matches the model in panel b. The y-scale in panels a and c was re-calibrated to correct for differences in the amount of sample and number of acquired scans (see Methods).

### NMR Analysis of Changes in the Fibrils’ Fuzzy Coat

The CP spectra lacked signals from the most flexible parts of the fibril, such as the fuzzy coat where the MW8 epitope is expected to reside. MAS ssNMR experiments based on insensitive nuclei enhanced by polarization transfer (INEPT)-type experiments allow observation of the exposed fuzzy coat. These experiments selectively highlight signals from mobile residues while filtering out the contributions from immobilized ones (Figure 3c).^[55]^ Consequently, the INEPT spectra revealed signals from the mobile flanking domains (Figure 3b, dark/light blue region) with the intensity determined by both the flexibility and the quantity of atoms detected. The spectra of the control fibrils matched those reported previously.^[15,30,31]^ Interestingly, the spectra measured in the presence of MW8 showed notable spectral changes: some peaks shifted, others disappeared, and some were reduced in intensity (Figure 3c, boxes 1 and 2). However, due to peak overlap in the 1D spectra, precise analysis of affected peaks was challenging. To address this, an INEPT-based 2D ^1^H-^13^C experiment was performed (Figure 3d), which provided peaks for a carbon and its directly bonded hydrogens, enabling a more precise assignment and comparison to previously assigned spectra (Figure S9).^[29]^ Observed chemical shifts are tabulated in Tables S1 and S2. The antibody binding predominantly affected specific residues. Peaks from leucine (L) and glutamate (E) disappeared from the INEPT spectrum (Figure 3c, box 4). Notably, these residues match the core residues of the MW8 epitope, in the tail of the PRD, based on the sequence AEEPLHRP. Additionally, a decreased proline peak intensity was observed. These intensity losses in INEPT spectra suggest reduced mobility of the impacted residues. Other residues, such as lysine (K), arginine (R), and histidine (H), exhibited a difference in chemical shift (Figure 3c, boxes 3 and 5, respectively). This would imply a retention of dynamics but a change in the (time-averaged) local structure and/or chemical environment. Here it is important to note that the His signals likely stem (in part) from the C-terminal His tag, which is known to experience polymorph-dependent motion similar to the C-terminal HTTex1 tail segment. ^[31]^

Thus, despite the MW8 antibody being present at a sub-stoichiometric amount, we observed by ssNMR a specific immobilizing effect of the antibody on its recognized epitope in the fibrillar assemblies. A more in-depth NMR analysis was challenging due to the small size of the sample, which was limited by the amount of available MW8 antibodies. It is worth noting that dynamic differences in the HTTex1 C-terminus are a hallmark of fibril polymorphism, and particularly of fibril bundling.^[15,31,56]^ As noted above, the MW8 also seemed to induce increased bundling, as seen by TEM, which may contribute to some of the effects on PRD domain mobility (more on this below).

### Observed Changes in Fuzzy Coat Order and Disorder

Next, we had a closer look at the differences in mobility between the control and antibody-treated samples. The intensities of the CP and INEPT signals were compared against the direct ^13^C excitation (DE) intensities.^[55,57–60]^ In the DE experiment, contrary to INEPT and CP, the intensities are related to the number of equivalent atoms, and this experiment can be considered semi-quantitative. The proline C_γ,_ C_β,_ and C_δ_ peaks (~27, ~30, and ~50 ppm) exhibited reduced intensity in the CP spectrum (Figure 3a, red arrows). This finding implies that the antibodies modulated the proline-rich parts of the PRD that flank the polyQ core, causing them to become less ordered and thus displaying reduced CP transfer. Interestingly, we also see a reduction in prominent proline peaks (~27, ~30, and ~50 ppm) in the INEPT-based spectrum (Figure 3c; red arrows). This would suggest an *increase* in proline ordering, leading to faster ^1^H and ^13^C T_2_ relaxation compromising the effectiveness of the refocused INEPT experiment.^[58–60]^ This seemingly contradictory behavior can be explained by the fact that the CP and INEPT analyses probe different parts of the PRD: CP monitors immobilized parts near the fibril core, while the INEPT shows signals from Pro residues in the more flexible tail end of the C-terminus. Therefore, INEPT signal reductions imply an ordering effect on the normally flexible tail of the PRD (i.e., near the MW8 epitope), while the reductions in the CP suggest an increased disorder in the rigid PRD close to the fibril core. In total, we observed that the antibody interaction enhances the rigidity of the PRD near the MW8 epitope while also reducing the structural order of polyproline regions near the rigid core.

### Protofilament Core Stability and Disruption

The above data point to a disordering or disruption of core-proximal flanking regions upon MW8 binding. This effect bore a resemblance to previous reports suggesting that chaperones may disaggregate fibrils through their interactions with amyloid flanking domains.^[61–63]^ To examine this possibility, we submitted the MW8-decorated fibrils to an extended incubation period and remeasured the fibril samples’ NMR signals. Doing so, we did not observe evidence of changes in spectra (Figure S10). If the fibrils were disaggregated or fundamentally remodeled by the antibodies, then one would have expected to see notable changes in the fibrils’ ssNMR spectra.^[31]^ Thus, it appears that MW8 can remodel or at least dis-order the fuzzy coat but not disrupt the stable core of these HTTex1 fibrils. Notably, the NMR was done on Q32-HTTex1 fibrils, which possess a smaller core than the aggregated typically found in HD patients. It is important to note that the ssNMR probes the integrity of the core of individual protofilaments (Figure 3a), which does not appear to change. However, it is separate from changes in the *supramolecular* fibril architecture may also be of significance and relevance (see below).

### Implications For HD Biology

We have observed by ssNMR that the MW8 antibody has a clear impact on the fuzzy coat of HTTex1 fibrils without disrupting the polyQ core itself. Interestingly, the TEM that probes the fibrils on a more architectural level, provided insights into supramolecular changes in how individual fibrils interact with each other. We noted that MW8 favored the bundling of fibrils into larger supramolecular assemblies. Prior work has suggested that small size and reduced stability may predict fibril-induced toxicity in the cellular environment.^[31,53,54,64,65]^ Arrasate and colleagues^[53]^ showed that the formation of larger fibril clusters may provide a protective effect by sequestering the toxic soluble species and converting them into less harmful aggregates. The reasons for isolated fibrils being potentially more toxic than large fibril clusters are diverse and multifaceted. Aberrant interactions of aggregated and misfolded proteins with cellular components are, by necessity, mediated by exposed segments and residues, with limited roles for buried (parts of) proteins. Moreover, it has been suggested that fibril surfaces (when exposed) may act as catalysts of toxic oligomer formation or as catalysts of the nucleation of new fibril species (via secondary nucleation type processes^[12,17,66,67]^). Moreover, the spreading of (auto-catalytic) amyloid entities between cells and cellular uptake is hindered by the formation of large inclusions. Cells can thus reduce the toxic consequences of protein misfolding and aggregation by sequestering proteins into restricted and tightly packed inclusions. Here, we would note that this does not necessarily imply that the relevant toxic species is necessarily non-fibrillar, but rather can be small and isolated fibrils. Such an idea is consistent with prior super-resolution microscopy studies as well as measurements of seeding-capable aggregates.^[64,65,68]^ Thus, an interesting implication of our MW8 studies is that we observed the increased clustering of HTTex1 fibrils. As illustrated in Figure S11, this could be rationalized by the multivalent binding by antibodies to multiple fibrils. This could offer a mechanism for increased clustering of fibrils, with the potential of reducing their toxic effects in cells. Thus, one favorable effect of fibril-binding antibodies may be not to disrupt or destroy them but rather to modulate the supramolecular morphology in ways that can reduce cellular interactions and cytotoxic effects.

## Conclusion

This study showed how ssNMR and EM could be utilized to probe the interaction between antibodies and HTTex1 fibrils. We observed that these interactions not only modified the mobility of the specific epitope but also affected other aspects of the fibril structure. Notably, the bound antibody disrupted the fuzzy coat regions near the epitope while simultaneously promoting enhanced fibril bundling. Instead of leading to a destabilization of the fibrils, this bundling could inhibit interactions with other proteins or cellular components, potentially impacting the mechanisms through which HTTex1 fibrils exert stress on and contribute to neuronal cell death. Our study highlights the potential role of ssNMR in analyzing the structural preferences of widely used HTTex1 and polyQ-binding antibodies, as well as amyloid-binding antibodies more broadly. This approach offers a unique chance to elucidate the structural and dynamic properties of bound substrates without requiring long-range order or crystallinity. Gaining deeper insights into these amyloid-antibody complexes is pivotal for accurately interpreting the widespread use of presumed structure-specific antibodies that are extensively employed in HD research and the amyloid field more generally.

## Supporting Information

The authors have cited additional references within the Supporting Information.^[69–75]^

## Acknowledgements

The authors would like to express their appreciation to Marc Stuart, the manager of the Electron Microscopy facility, for his guidance and training, the Developmental Studies Hybridoma Bank, created by the NICHD of the NIH and maintained at The University of Iowa, Department of Biology, Iowa City, IA 52242, and the CHDI foundation for providing antibodies. This research was supported by funds from the University of Groningen, the CampagneTeam Huntington foundation, and the European Huntington’s Disease Network (EHDN; seed fund project 1185).

