## Supporting Information for "Probing How Anti-Huntingtin Antibodies Bind the Fibrillar Fuzzy Coat Using Solid-State NMR"

### 1. Experimental section

#### 1.1 General information

Monoclonal antibodies (MW1, MW7, and MW8), developed by Ko and coworkers, were obtained from the Developmental Studies Hybridoma Bank (DSHB), created by the NICHD of the NIH and maintained at The University of Iowa, Department of Biology, Iowa City, IA 52242. The antibody 3B5H10 was obtained from Sigma Aldrich, and 2B7 from the Coriell Institute (cat# CH02024). Q32- and Q44-HTTex1 were generated with a terminal HIS-tag and as fusion proteins with N-terminal maltose-binding protein (MBP) solubility tags, as the following constructs: MBP-Q32-HTTex1 and MBP-Q44-HTTex1. These constructs were cloned into pMALc5x plasmid (Q32) and pMALc2x plasmid (Q44), by Genscript (Piscataway, NJ).[1]

#### 1.2 Protein production

The fusion proteins were overexpressed in E. coli BL21(DE3) pLysS cells (Invitrogen, Grand Island, NY). Cells were cultured in LB medium supplemented with ampicillin and chloramphenicol at 37°C with agitation at 200 rpm, until reaching an optical density (OD 600) of 0.8. Subsequently, the temperature was gradually reduced from 37°C to 18°C over 30 minutes while maintaining agitation at 200 rpm in preparation for induction. Uniform labeling of proteins with $^{13}C$ and $^{15}N$ isotopes was achieved using M9 minimal media supplemented with $^{13}C$-D-glucose and $^{15}N$ ammonium chloride (Sigma-Aldrich) during overexpression, as outlined in prior studies.[2] Protein production was induced by adding 0.6 mM IPTG (Sigma-Aldrich), followed by 16 hours of expression at 18°C. Harvesting involved pelleting the cells at 5250×g for 20 minutes, resuspending them in phosphate-buffered saline (PBS; 10 mM sodium phosphate dibasic, 1.8 mM potassium phosphate monobasic, 137 mM sodium chloride, and 2.7 mM potassium chloride, pH 7.4), and cooling on ice. To this, 1 mM phenylmethanesulfonyl fluoride and 0.5 mg/mL lysozyme (both from Sigma-Aldrich) were added. Cell lysis was achieved via sonication (VCX130 Vibra-Cell Sonicator, Sonics & Materials, Inc) at 70% amplitude for 10 minutes, alternating 10-second pulses with 10-second pauses. Debris was cleared by centrifugation at 125,000×g for 45 minutes using a Beckman Coulter Optima LE-80K ultracentrifuge, and the soluble protein fraction was filtered through 0.45 µm sterile syringe filters (Fisherbrand – 15216869). Purification was carried out using a HisTrap HP nickel column (GE Healthcare) with an imidazole gradient, followed by dialysis into imidazole-free PBS using a 12,400 MWCO membrane (seamless high retention cellulose tubing, Sigma-Aldrich - D0405). Protein concentration was determined by measuring absorbance at 280 nm (average of three readings) using a JASCO V-750 spectrophotometer and a 10 mm quartz cuvette. The extinction coefficient, calculated using the ProtParam tool from ExPASy, was estimated at 66,350 $M^{-1}cm^{-1}$.[3] Protein purity, isotopic enrichment, and molecular weight were confirmed using ESI-TOF MS and SDS-PAGE (12%).[1]

#### 1.3 Fibril production

Cleavage of the purified fusion protein was carried out using Factor Xa (FXa; Promega, Madison, WI) at room temperature, in a 400:1 molar ratio (HTTex1:FXa), to release the HTTex1. SDS-PAGE (Bio-Rad Mini-Protein Precast TGX Gels, 12%) was used to monitor the cleavage. The aggregation process was allowed to proceed for 96 hours. The resulting HTTex1 fibrils were separated by centrifugation at 3000×g for 20 minutes, using a Thermo Scientific SL40R large bench centrifuge, and the supernatant was removed. The fibrils were washed at least three times with PBS buffer. For magic angle spinning (MAS) solid-state NMR (ssNMR) experiments, fibrils were incubated overnight with MW8 antibodies at a 4:1 molar ratio (HTTex1 monomer equivalent: antibody) to allow for binding. The supernatant's absorbance was measured in a 50 µl quartz cuvette with a 10 mm path length by a JASCO V-750 spectrophotometer, and the MW8 epsilon was calculated with ExPASy,[3] to confirm the absence of unbound antibodies. Prior to further analysis, the fibrils were pelleted and thoroughly washed to eliminate any remaining unbound antibodies.

1.4 Dot blot assay

A dot blot assay was performed to evaluate the interaction between the antibody and both the fused protein (used as monomer) and the aggregated HTTex1 fibrils, using a Bio-Dot apparatus (BIO-RAD). 1μg of fibril or fused protein was placed on a pre-wetted nitrocellulose membrane in Tris-buffered Tris buffer saline (TBS) (20 mM of Tris, 150 mM NaCl, pH 7.6) and allowed to slowly filter through. The apparatus was then washed with 100 μL of TBST (TBS with 0.1% v/v Tween 20 detergent, pH 8) solution, to ensure that all the protein passes through the membrane. Each well was subsequently washed three times with 300 μL of TBST. The nitrocellulose membrane was removed, placed, and incubated in blocking buffer (1% w/v BSA in 100 mL of TBST) for 1h at room temperature, followed by a wash with TBST. The membrane was then incubated with the primary antibody overnight at 4° C, using the following dilutions: 1:200 for MW1 and MW7,[4] 1:20000 for 2B7, 1:5000 for 3B5H10, [5] and 1:1000 for MW8.[6] After washing with TBST to remove unbound antibodies, the membrane was incubated with secondary antibodies (1:10000 dilution of Goat anti-Mouse IgG/IgM (H+L) Secondary Antibody, Alexa Flour 488 conjugate from Thermo Fisher) at r.t. for 1h. The membrane was then air dried and visualized LAS-3000 (FUJI FILM) imager system with a 460 nm light source and a 4515-DI filter. Data analysis was performed using the ImageJ software.[7]

1.5 Transmission electron microscopy (TEM)

Negative stain TEM was employed to analyze the sample. All samples were suspended in Milli-Q water and deposited onto plain carbon support films mounted on 200-mesh copper grids (SKU FCF200-Cu-50, Electron Microscopy Sciences, Hatfield, PA). The final concentration of 40 μM for the fibrils with and without antibodies and to 25 μM for the MW8 alone. The grids were glow-discharged for 30 seconds to 1 minute, before sample deposition. Excess Milli-Q water was blotted off, and fibrils were negatively stained with a 2% (w/v) solution of uranyl acetate, applied immediately after blotting, for 30 seconds to 1 minute. Excess stain was removed by blotting, and the grids were air-dried. Imaging was performed using a Philips CM120 electron microscope operating at 120 kV. Images were captured on a slow-scan CCD camera (Gatan). Fibril widths were measured transversely to the fiber long axis using the straight free-hand tool in Fiji (ImageJ).[7] Measurements spanned the negatively stained portions of the fibrils; observed width is assumed to reflect the polyQ core, with the stain accumulating between the flanking domains. In low-resolution images, diameters were measured in regions with the clearest boundaries, with three measurements taken per fibril unless the width varied significantly. In specific cases, width measurements were validated on isolated fibrils using Fiji's Plot Profile tool, which provides average grayscale intensity profiles across the fibril axis. For comparison of observed fibril and antibody sizes, the antibody structural analysis was performed on ChimeraX using the structure of a similar IgG2a antibody (PDB code: 5DK3).[8]

1.6 Solid-state NMR experiments (ssNMR)

The prepared material was packed into 3.2 mm zirconia regular-wall MAS rotors (Bruker Biospin, Billerica, MA) by pelleting the hydrated sample directly into the rotor. Pelleting was achieved using an ultracentrifuge packing device, at ~130,000 g in a Beckman Coulter Optima LE-80K ultracentrifuge equipped with an SW-32 Ti rotor.[9] Before sealing the rotor, the excess water was removed, and an insert was placed between the sample and the drive cap. The wet sample weight (including proteins and buffer) for each rotor was ~16 mg for the fibrils with MW8 and ~18 mg for the control, with the former containing ~0.1 mg of HTTex1 fibrils and the latter ~3.7 mg of HTTex1 fibrils. Solid-state NMR studies were conducted on the hydrated, unfrozen samples under magic angle spinning (MAS) at a rate of 13 and 10 kHz. Spectra were collected on a Bruker AVANCE NEO 600 MHz spectrometer with a 3.2 mm Bruker HCN MAS probe equipped with an HCN Efree coil. The NMR measurements were performed with the temperature control system configured for a setpoint temperature of 275K. The actual sample temperature will be higher due to frictional heating associated with the MAS.

The 1D $^{13}$C spectra were recorded using the following conditions: $^{1}$H 90° pulse was set to 3 μs, corresponding to a rf power of ~83 kHz, and $^{13}$C 90° pulse was set to 5 μs, corresponding to a rf power of ~50 kHz for the Cross Polarization (CP) experiment. For the Insensitive Nuclei Enhanced by

Polarization Transfer (INEPT) and Direct Excitation (DE), the $^{1}H$ 90° pulse was set to 3 μs corresponding to a rf power of ~83 kHz, and the $^{13}C$ 90° pulse was set to 4 μs, corresponding to a rf of 60 kHz. The CP step used a 70−100% ramped-amplitude (RAMP) shape on the $^{1}H$ channel and a square-shaped pulse on the $^{13}C$ channel at 50 kHz. During data acquisition, the TPPM (two-pulse phase-modulated) decoupling scheme was applied, with a basic decoupling unit pulse length of 5.8 μs and an RF field strength of ~83 kHz. The 2D $^{13}C$-$^{13}C$ INEPT-HETCOR experiments were recorded setting the $^{1}H$ 90° pulse and $^{13}C$ 90° at 4 μs corresponding to a rf power of ~60 kHz, and $^{1}H$ decoupling during acquisition time of ~71.4 kHz.

The chemical shifts for $^{13}C$ ssNMR were indirectly referenced to liquid ammonia and aqueous DSS, respectively, by measuring adamantane $^{13}C$ signals.[10] Spectra were processed using Bruker Topspin and NMRPipe,[11] using (exponential) line broadening of 30 to 40 Hz, as indicated below. The spectra were visualized using Topspin and CcpNmr software 2.5.[12] The 1D $^{13}C$ DE experiments were deconvoluted in Dmfit, applying mixed Gaussian-Lorentzian functions.[13] To permit visual inspection of the 1D spectra for the control and antibody-containing samples, we applied a vertical scaling correction to account for the difference in protein amount (in terms of the $^{13}C$-labeled HTTex1 protein) and the number of acquired scans.

### 2. Supplementary data

Table S1. $^{13}C$ chemical shifts of assigned residues in the HTTex1 control sample. The standard deviation of the $^{13}C$ chemical shifts is ± 0.1-0.3 ppm based on the systematic comparison, during the peak assignment process, of peak positions between 1D and 2D spectra. In parentheses, the chemical shift of peaks that changed upon antibody binding is indicated, in case of peak disappearance, not available (N/A) is reported. Peak assignments were based on prior reports.[14] See also Figure S7 for the 2D spectra with assignments.

| | **C'** | **Cα** | **Cβ** | **Cγ** | **Cδ** | **Cε** | **Visible by CP** | **Visible by INEPT** |
|---|---|---|---|---|---|---|---|---|
| Met (M) | | | | | | 17.2 | | + |
| Ala (A) | | 52.8 | 19.5 | | | | | + |
| Thr (T) | | | | 20.7 | | | | + |
| Leu (L) | | 55.7(N/A) | 42.5 (N/A) | 27.2 | 25.3; 23.8 | | | + |
| Glu (E) | | 57.7(N/A) | | 36.4(N/A) | | | | + |
| Lys(K) | | | | 25 (24.4) | | 43.7 | | + |
| Phe (F) | | 58.8 | | | | | | + |
| Ser (S) | | 64.1 (64.2) | | | | | | + |
| Gln ‘a’ (a:Q) | 176.1 | 56 | 34.4 | 34.4 | 178.8 | | + | |
| Gln ‘b’ (b:Q) | 174.4 | 54.2 | 32 | 30.8 | 177.9 | | + | |
| Pro PPII ($P_{PPII}$) | | 61.5 | 30.9 | 27.6 | 50.7 | | + | |
| Pro RC ($P_{RC}$) | | 63.3 | 32.4 | 27.6 | 50.7 | | + | |
| Gln (Q) | | 53.8 | 29.3 | 33.9 | | | | + |
| Gly (G) | | 45.5 | | | | | | + |
| Val (V) | | 62.2 (N/A) | | 21.4 | | | | + |
| His (H) | | 56.3 (55.5) | 30.9 (29.4) | | 119.5 | 138.5 | | + |
| Arg (R) | | 53.7 (55.2) | | | | | | + |

Table S2. $^{1}H$ chemical shifts of assigned residues in the dynamic fuzzy coat of our Q32-HTTex1 fibril samples. The standard deviation of the $^{1}H$ chemical shifts is ± 0.1-0.3 ppm based on the systematic comparison, during the peak assignment process, of peak positions 2D spectra. Reported values are for the control fibrils, without antibody; in parentheses, the chemical shift of peaks that changed upon antibody binding is indicated; in case of peak disappearance, N/A is reported. Peak assignments were based on prior work.[14] See also Figure S7 for the 2D spectra with assignments.

| **Amino Acid** | **Hα(ppm)** | **Hβ (ppm)** | **Hγ (ppm)** | **Hδ (ppm)** | **Hε (ppm)** |
|---|---|---|---|---|---|
| Met (M) | | | | | 2.1 |
| Ala (A) | 4.3 | 1.4 | | | |
| Thr (T) | | | 0.9 | | |
| Leu (L) | 4.3 (N/A) | 1.6 (N/A) | 1.6 | 0.9; 0.9 | |
| Glu (E) | 4.4 (N/A) | 1.6 | 2.3 (N/A) | | |
| Lys(K) | | | 1.9 (2) | | 3.2 |
| Phe (F) | 4.4 | | | | |
| Ser (S) | 3.9 (3.8) | | | | |
| Pro PPII ($P_{PPII}$) | 4.7 | 1.9; 2.4 | 2 | 3.7 | |
| Pro RC ($P_{RC}$) | 4.4 | 2.3; 1.9 | 2 | 3.7 | |
| Gln (Q) | 4.6 | 2.1; 2 | 2.4 | | |
| Gly (G) | 4 | | | | |
| Val (V) | 4.1 (N/A) | 2.1 (N/A) | 0.9 | | |
| His (H) | 4.6 (4.7) | 3 (3.2) | | 6.9 | 7.9 |
| Arg (R) | 4.6 | | | | |

Table S3. Experimental conditions of ssNMR experiments. Abbreviations: NS, number of scans per $t_1$ point; Temp., temperature; MAS, magic angle spinning rate; RD, recycle delay; TPPM, $^1$H decoupling power during evolution and acquisition using the two-pulse phase modulation scheme; $t_1$ evol., maximum $t_1$ evolution time expressed in number of $t_1$ points (real+imaginary) x $t_1$ increment time. All NMR experiments were performed using a setpoint for the NMR temperature control system of 275K.

| | **Sample** | **Experiment** | **NS** | **MAS (kHz)** | **RD (s)** | **TPPM (kHz)** | **CP contact time (ms)** | **$t_1$ evol. (μs)** |
|---|---|---|---|---|---|---|---|---|
| Figure 3a; S7 | Q32- HTTex1 control | $^{13}$C CP | 512 | 13 | 3 | 83.3 | 1 | N/A |
| Figure S7 | Q32- HTTex1 control | $^{13}$C DE | 512 | 13 | 3 | 83.3 | N/A | N/A |
| Figure 3c; S7 | Q32- HTTex1 control | $^{13}$C INEPT | 512 | 10 | 3 | 83.3 | N/A | N/A |
| Figure 3a; S8 | Q32- HTTex1 +MW8 | $^{13}$C CP | 56832 | 13 | 3 | 83.3 | 1 | N/A |
| Figure S8 | Q32- HTTex1 +MW8 | $^{13}$C DE | 56832 | 13 | 3 | 83.3 | N/A | N/A |
| Figure 3c; S8 | Q32- HTTex1 +MW8 | $^{13}$C INEPT | 56832 | 10 | 2.8 | 83.3 | N/A | N/A |
| Figure 3d; S9 | Q32- HTTex1 control | $^1$H - $^{13}$C INEPT HETCOR | 128 | 10 | 2.8 | 83.3 | N/A | 1636x 151.48 |
| Figure 3d; S9 | Q32- HTTex1 +MW8 | $^1$H - $^{13}$C INEPT HETCOR | 768 | 10 | 2.8 | 71.4 | N/A | 1636x 151.48 |
| Figure S10 | Q32-HTTex1 +MW8 after long incubation | $^{13}$C INEPT | 512 | 10 | 3 | 83.3 | N/A | N/A |

#### 3. Supplementary images

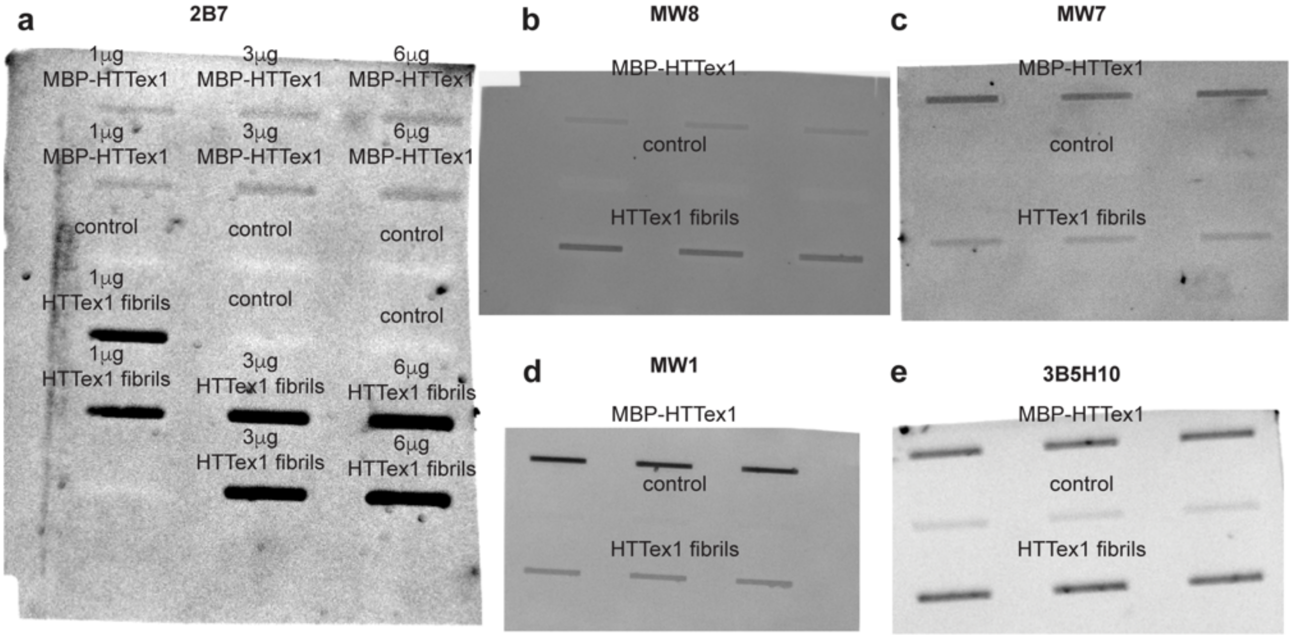

*Figure S1. Dot blot assay of Q32-*HTTex1 *fused protein and fibrillar form with different types of antibodies. a) interaction with 2B7 antibody with different amounts of fused protein (1, 3, and 6 μg) and fibril (1, 3, and 6 μg). b) interaction with MW8 antibody. c) interaction with MW7 antibody. d) interaction with MW1 antibody. e) interaction with 3B5H10 antibody. The antibodies 2B7 and 3B5H10 are widely used antibodies with affinities for the HTT$^{NT}$ and polyQ segments of HTTex1, and they are included here as additional biochemical characterization of the fibril polymorphs used in this study.*

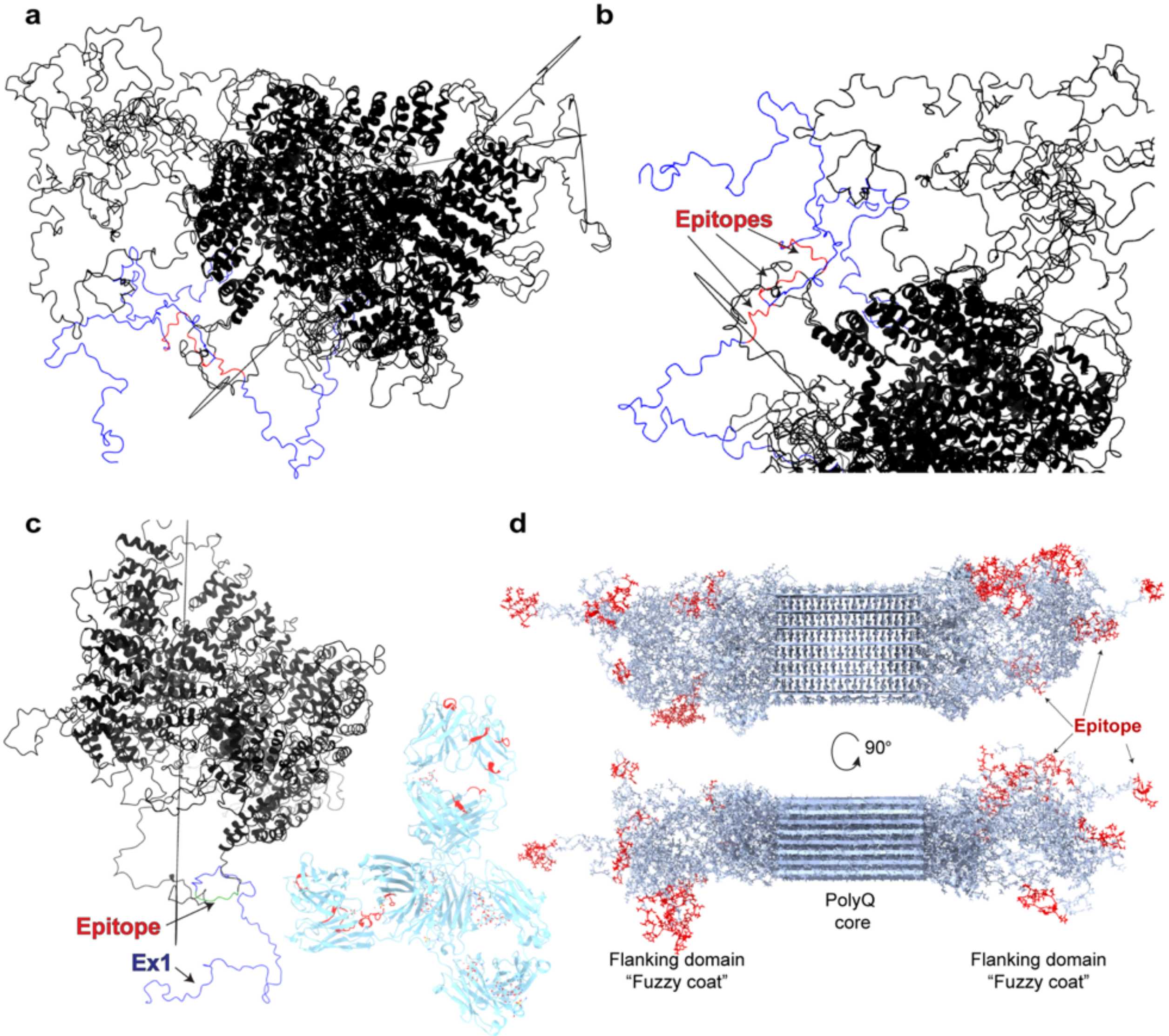

*Figure S2. 3D model of the wild-type Q23-HTT protein with HAP40 bound, shown in black. a) 3D model of the wild-type Q23 HTT, including the intrinsically disordered region (IDR), simulated using 3D modeling, such as HTTex1. Due to the high mobility of the IDR, those regions are highly challenging to study and, for this reason, are simulated, offering different conformations. Three possible models of HTTex1 regions are colored in blue, and the MW8 epitope in red. b) Zoomed view of MW8 epitope in the HTTex1 regions to illustrate how the epitopes are close to another HTT domain, making it inaccessible for MW8 recognition. c) 3D structures of wild-type Q23-HTT protein in black, with HTTex1 in blue and MW8 epitope in green; and IgG2a antibody in cyan, with in red the binding regions. It illustrates how the full-length protein impairs antibody binding due to steric hindrance. d) Top and side view of HTTex1 fibrils in gray with the MW8 epitope in red. As visible, the epitope is present only in the "fuzzy coat" and not in the core. The PDB file of Q23 HTT is from reference [15], the antibody has the PDB code 5DK3, and the HTTex1 model is from reference [16]. Both models were analyzed using ChimeraX.[17] The highlighted regions were selected using the selection sequence tool in ChimeraX.*

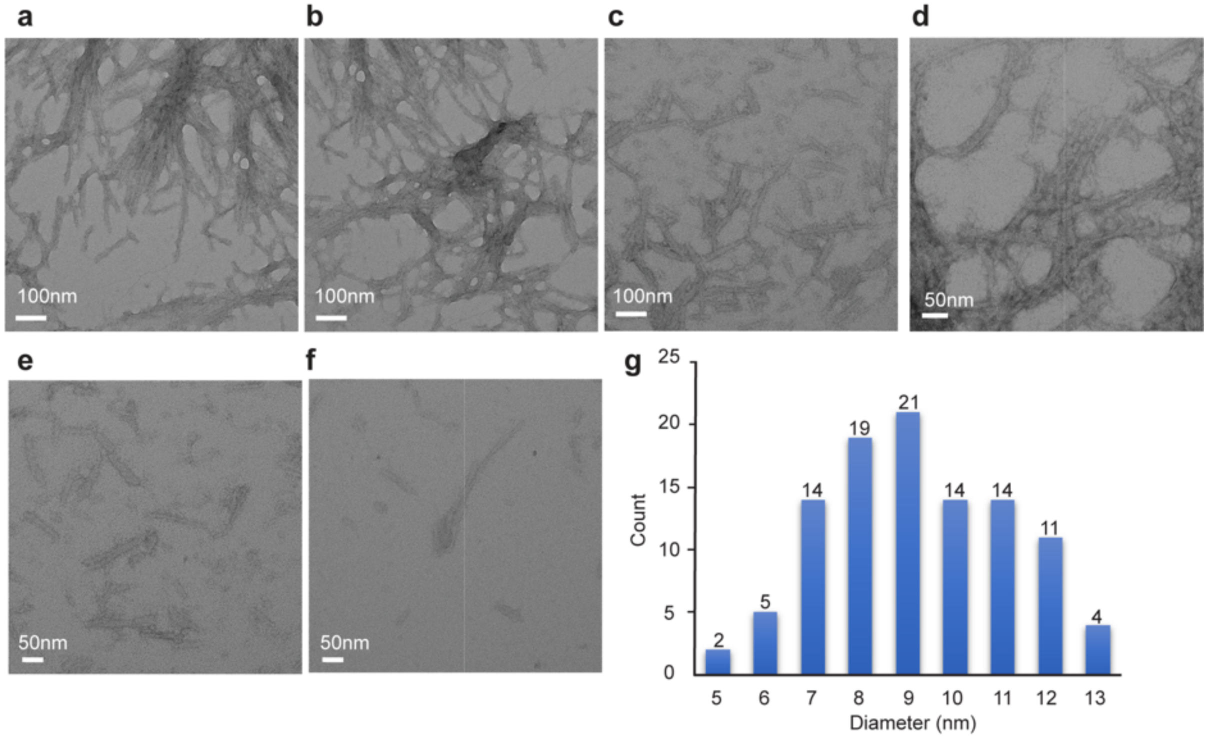

*Figure S3. Negative stain TEM images of Q44-HTTex1 fibrils. a,b,c) TEM data at the magnification level of 45000x. d,e,f) TEM data at the magnification level of 60000x. g) Fiber width distribution of Q44-HTTex1 fibrils aggregated at 25 °C and measured with the tool plot profile of Image J.[7]*

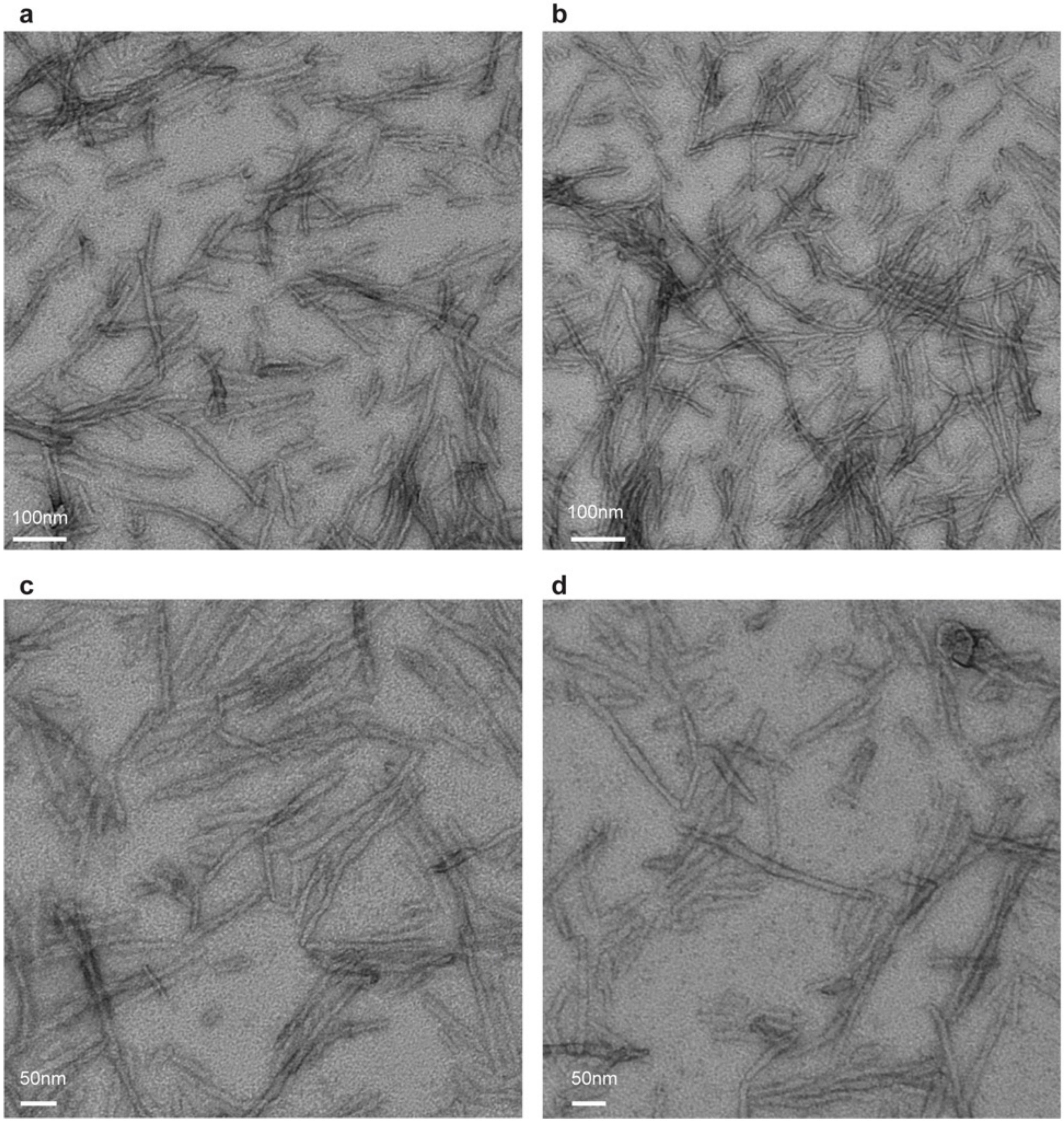

*Figure S4. Negative stain TEM images of Q32-HTTex1 fibrils. a,b) TEM data at the magnification level of 45000x. c,d) TEM data at the magnification level of 60000x.*

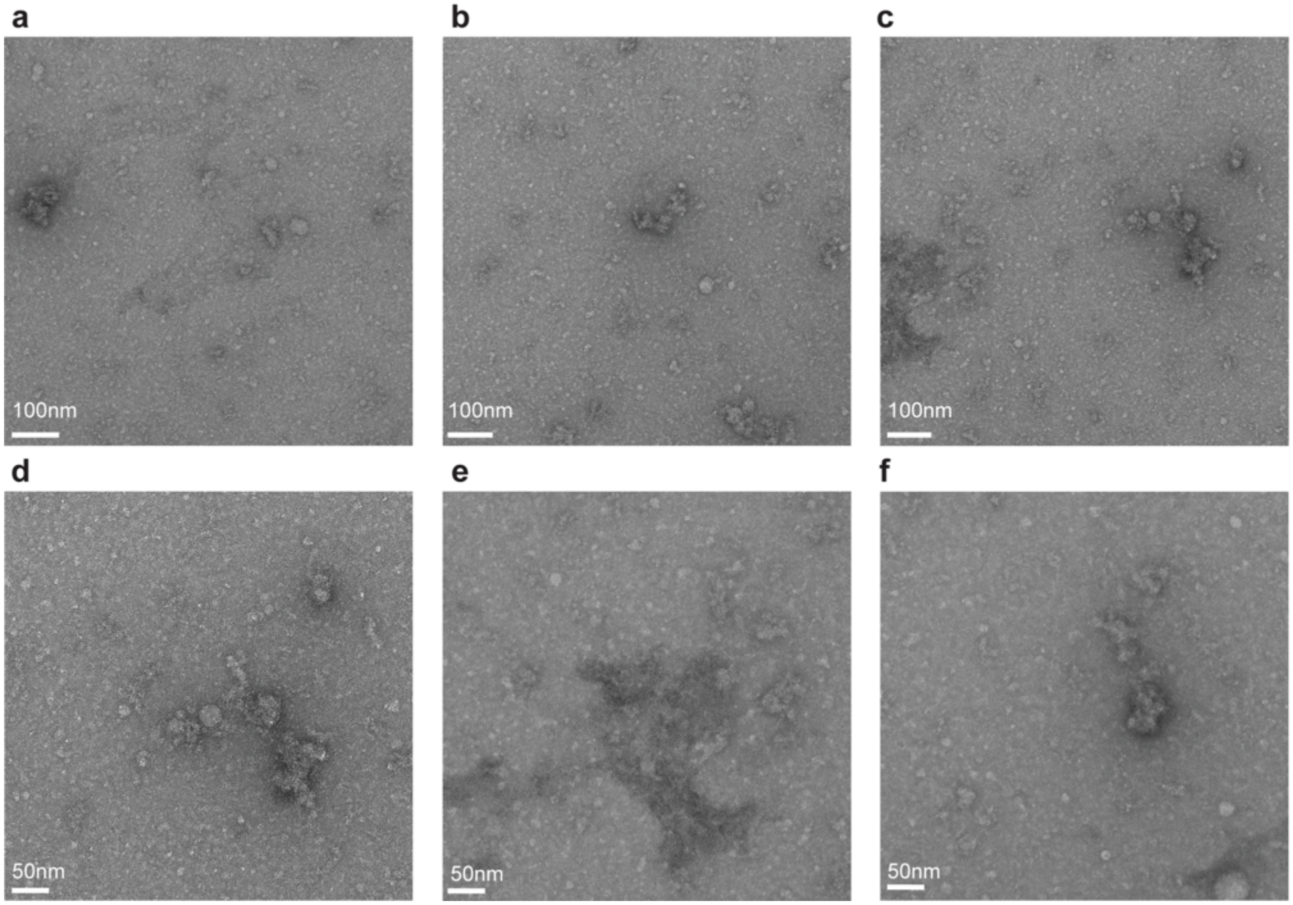

*Figure S5. Negative stain TEM images of MW8. a,b,c) TEM data at the magnification level of 45000x. c,d,e) TEM data at the magnification level of 60000x.*

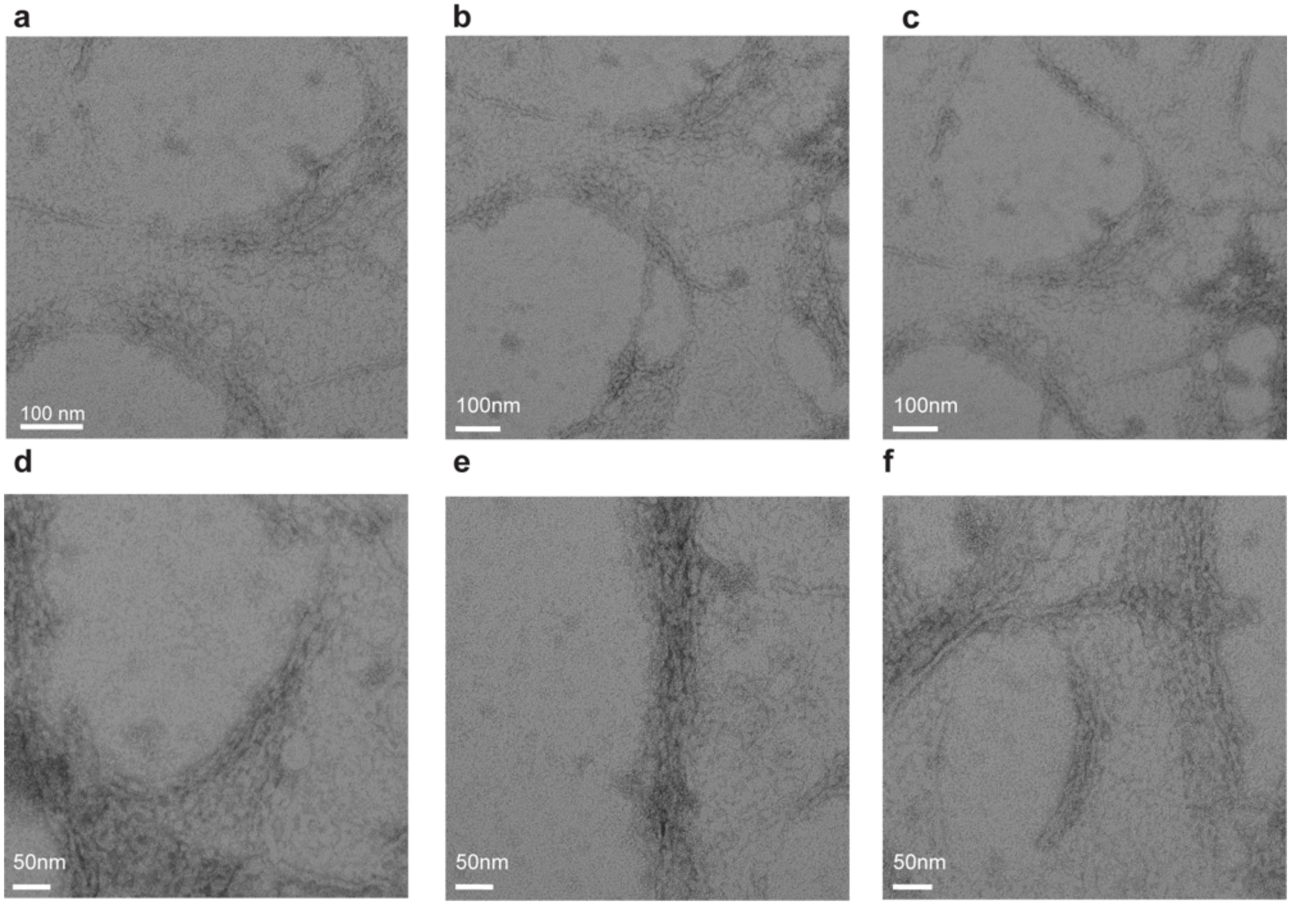

*Figure S6. Negative stain TEM images of MW8 bound to Q44-HTTex1. a,b,c) TEM data at the magnification level of 45000x. c,d,e) TEM data at the magnification level of 60000x.*

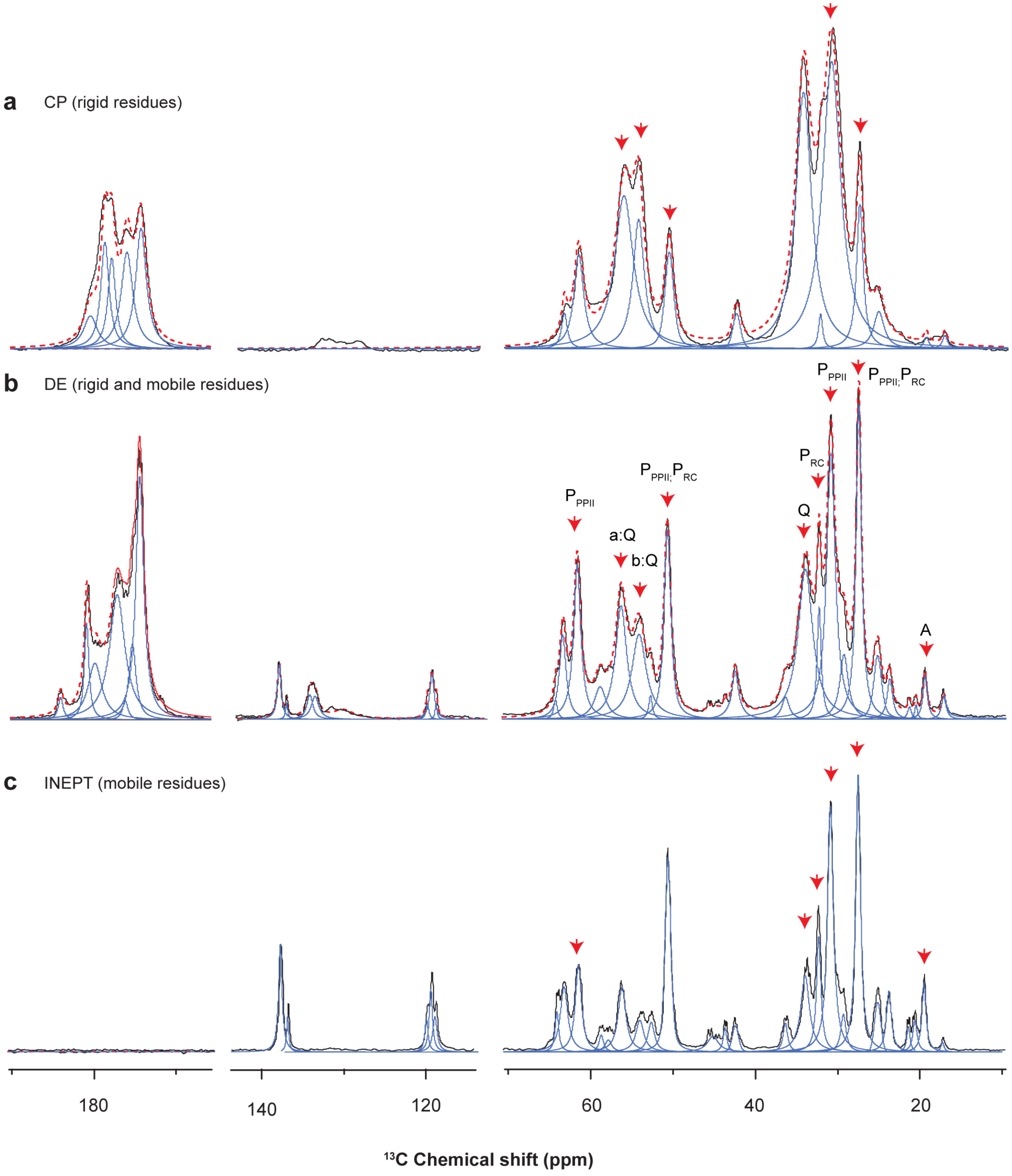

Figure S7. 1D $^{13}$C spectra of the HTTex1 control sample. a) $^{13}$C cross-polarization (CP) spectrum showing the rigid residues of the HTTex1 fibrils. b) $^{13}$C direct excitation (DE) spectrum showing the rigid and mobile residues of the HTTex1 fibrils. c) $^{13}$C insensitive nuclei enhanced by polarization transfer (INEPT) spectrum showing the mobile residues of the HTTex1 fibrils. The measured spectrum is shown as a black solid line, deconvolved peaks as a blue solid line, and the summed model as a red dashed line. The aliphatic regions of the CP and INEPT spectra are also represented in the main text in Figures 3a and c. The spectra were processed with Topspin using an exponential line broadening of 40 Hz, and further analyzed in the Dmfit program[13] (see Methods). Signals from the C-terminal His tag are visible in the aromatic region (110-145 ppm).

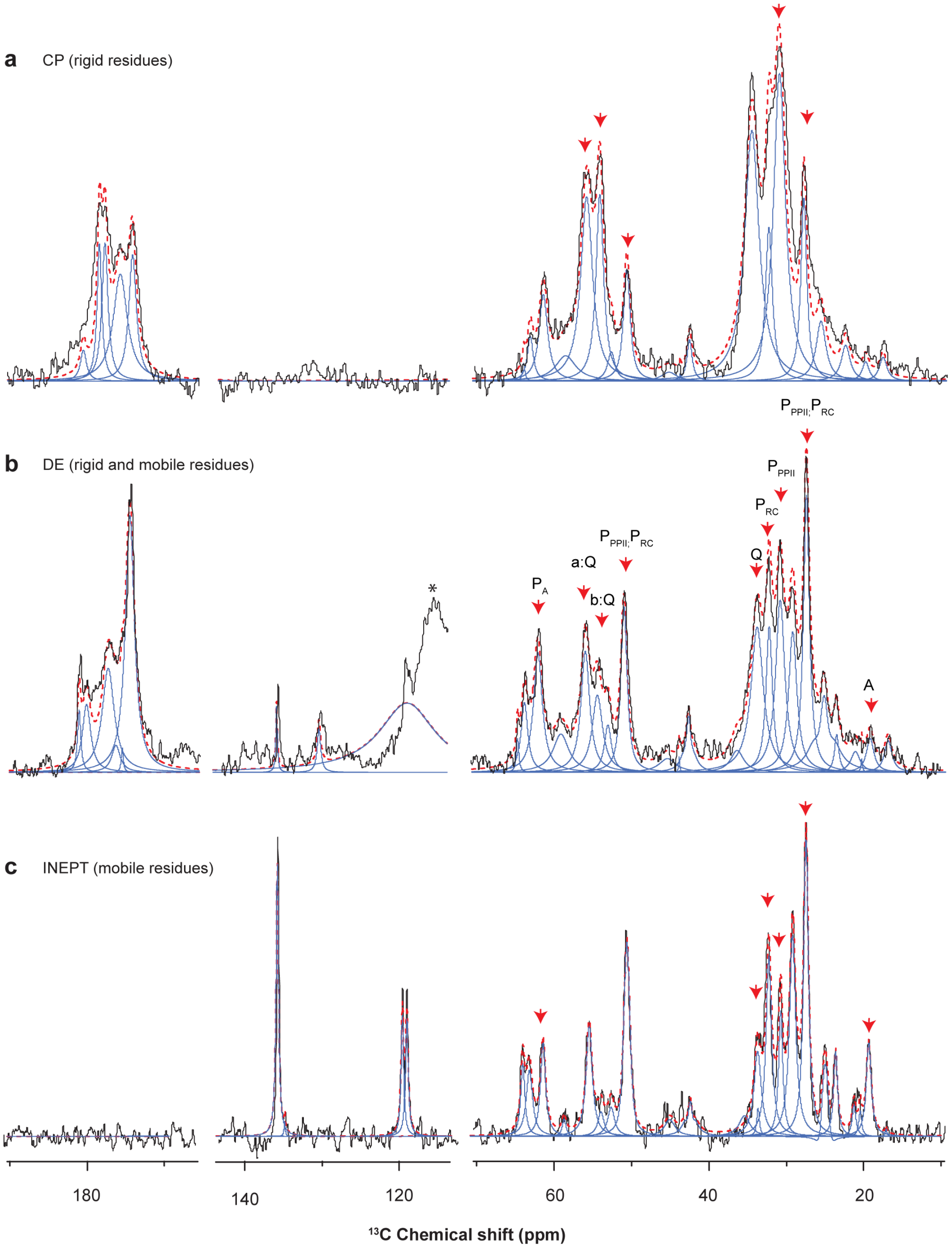

Figure S8. 1D $^{13}$C spectra of the sample HTTex1+MW8. a) $^{13}$C cross-polarization (CP) spectrum showing the rigid residues of the HTTex1 fibrils. b) $^{13}$C direct excitation (DE) spectrum showing the rigid and mobile residues of the HTTex1 fibrils. c) $^{13}$C insensitive nuclei enhanced by polarization transfer (INEPT) spectrum showing the mobile residues of the HTTex1 fibrils. The experimental spectrum is represented with a black solid line. The deconvolved peaks are shown as a blue solid line, while their combined sum shown as a red dashed line. The aliphatic regions of the CP and INEPT spectra are also shown in the main Figure 3. The limited signal:noise and peak overlap prevented a robust quantitative comparison between the samples. Signals from the C-terminal His tag are visible in the aromatic region (110-145 ppm), with partial overlap of a sample spacer (Kel-F) in the DE spectrum (marked with *).

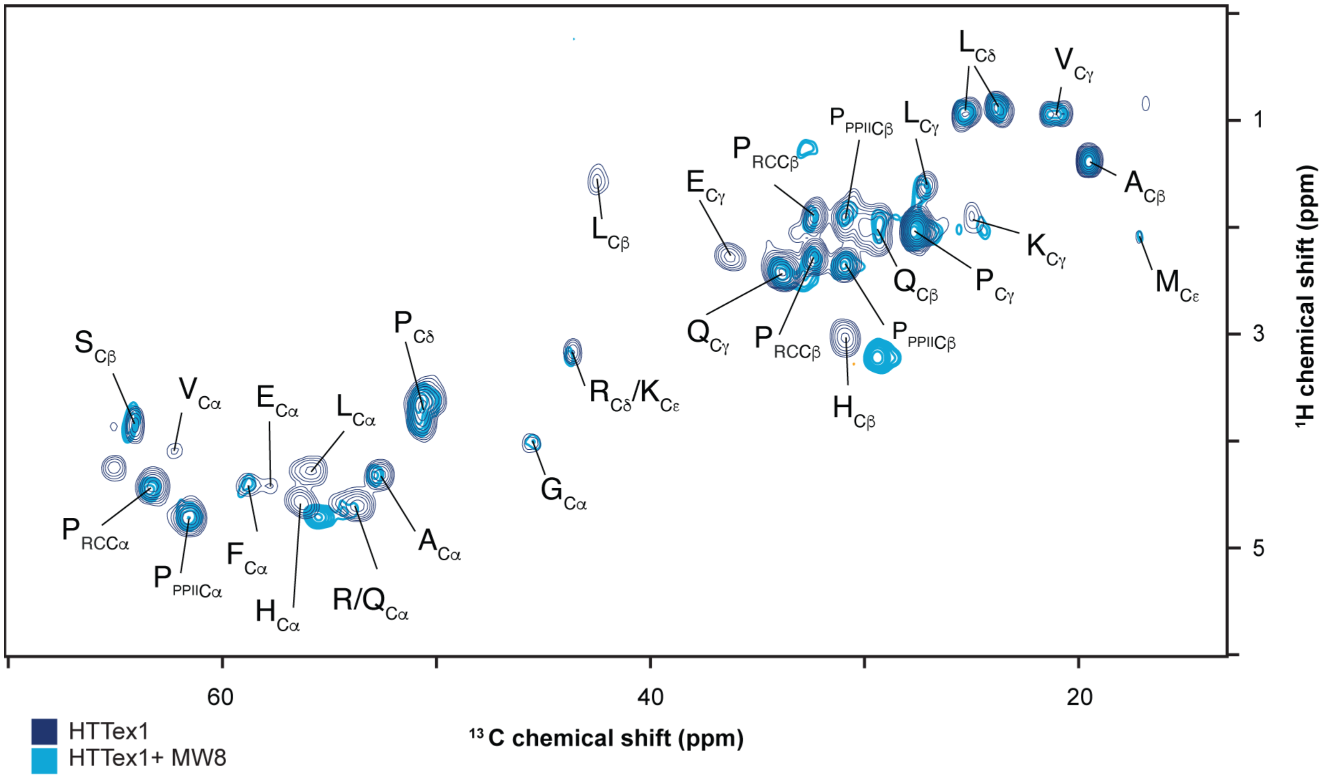

*Figure S9. 2D $^{13}C$ INEPT-HETCOR ssNMR spectra of the HTTex1 (dark blue) and HTTex1+MW8 (light blue) from the main text, indicating peaks labels according to residue type and aliphatic carbon type.*

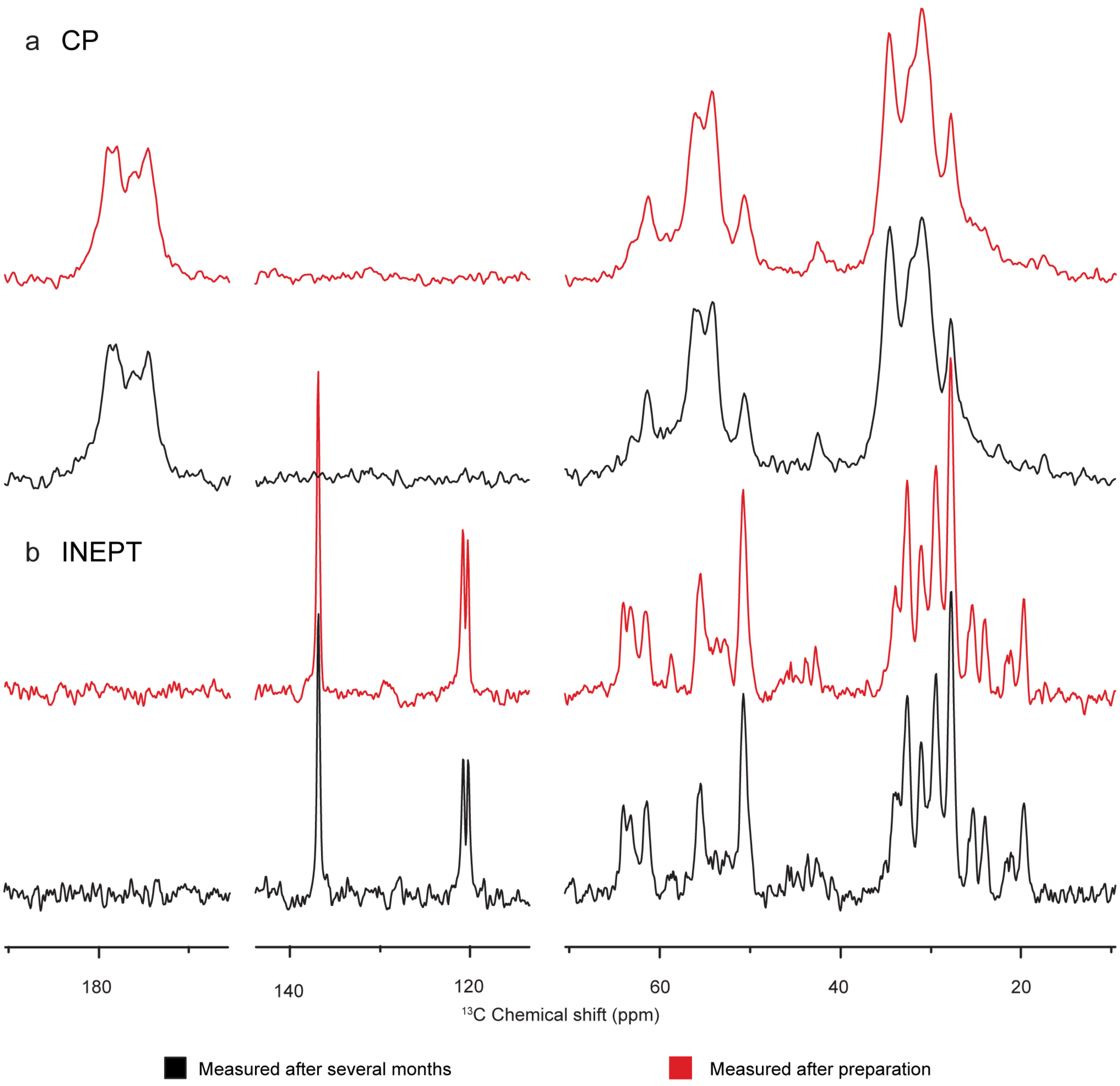

*Figure S10. 1D $^{13}C$ NMR spectra of HTTex1 with antibodies measured after several months of storage. a) Comparison of 1D $^{13}C$ NMR CP spectra and b) 1D $^{13}C$ NMR INEPT spectra of HTTex1+MW8 measured immediately after sample preparation (red spectrum) and several months later (black spectra). The processing was performed using an exponential line broadening of 30 Hz.*

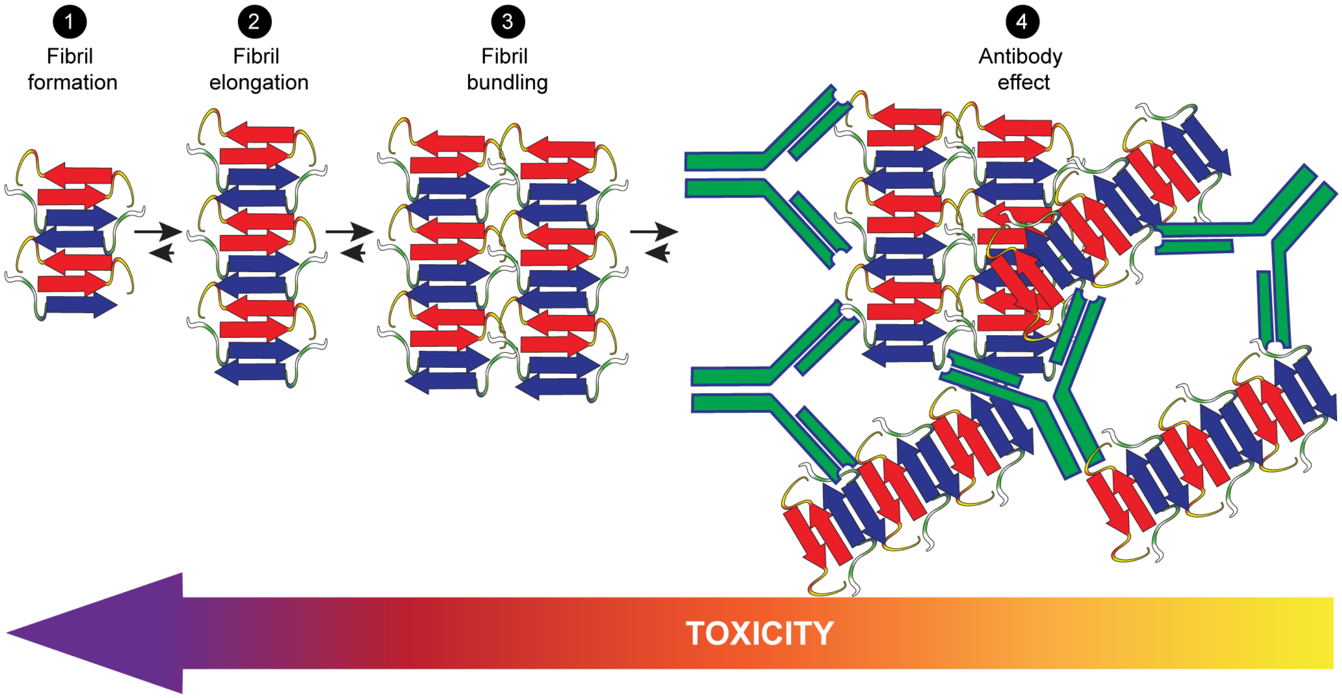

*Figure S11. Schematic representation of different levels of HTTex1 aggregation and fibril bundling. (1) Unfolded HTTex1 fragments undergo aggregation to form isolated fibrils. (2) Fibrils can elongate as additional monomers are incorporated. (3) In addition, fibrils can interact with one another via their fuzzy coat, manifesting as bundling. Bundling may occur for pre-formed fibrils but may also derive from secondary nucleation processes. In either case, interactions between non-polyQ flanking segments appear to be involved in bundling. (4) The presence of antibodies promotes further bundling, resulting in the formation of larger fibril clusters. The colored arrow at the bottom is meant to indicate that the toxicity of fibrils may vary as a function of the aggregate/cluster size. Larger clusters may be less toxic to cells compared to small fibrils, oligomers, or fibrillar fragments. The underlying mechanisms may be diverse (see main text).*
